# Effect of urbanisation on soil and plant pollution in province of Valencia, Spain)

**DOI:** 10.1101/2024.03.27.586674

**Authors:** Edina Simon, Ismail Haziq Bin, Bianka Sipos, Vanda Éva Abriha-Molnár, Dávid Abriha, Oscar Andreu-Sanchez, Rafael Boluda, Luis Roca-Pérez

**Author notes:** Corresponding author: E. Simon.

## Abstract

The accumulation of the toxic metals can result in the degradation of neighboring ecosystems and reducing their value to humans in terms of aesthetic properties and utility. Therefore, the aim of our research was to investigate the effect of urbanization on metal levels in soil samples and *Citrus sp*. tree leaves along an urbanization (rural, periurban and urban) in Valencia (Spain). There were significant differences for the content of clay, pH, electrical conductivity (EC), soil organic matter (SOM), concentration of Ca and exchangeable sodium percentage (ESP). A significant difference was not found in the case of elemental concentration of soil. There were significant differences for the concentration of Cu on *Citrus* sp. leaves, only. Based on our findings Valencia did not exhibit a significant relationship between the element concentrations and the urban gradient except for the case of copper in the leaf samples. The find no difference in the soil indicate the homogenous nature of the Valencia soil .

## Introduction

The urbanization is defined as the transition from an economy centered around agriculture products into a service economy with the use of technology and facilities to reduce the cost of production (Liang & Yang, 2019). Through the viewpoint of demography, urbanization is the aggregation of the population from rural communities into highly dense cities (Uttara et al. 2012, Kuddus et al. 2020). In the case of environmental sciences, the study of urbanization follows the impact of urban expansion and the urban city itself in regard to the changes that it incurs to the surrounding environment.

The main type of pollutant was classified as atmospheric pollutants which were sourced from vehicles and factory emissions in the context of an urban environment. For pollutant measurement in soil and air pollution was through the using of bioaccumulator or bioindicators which can absorb or collect pollutants into their systems (Brookes et al. 1995, LaGreca & Stutzman 2006).

A study by Rodríguez Martín et al. (2015) concluded that the increase in urban activities in the city of Valencia over a 70-year period had increased the levels of Cr, Ni and Cd in leaves of tree species of the Botanical Garden of Valencia.

Anthropogenic activities such as atmospheric emissions from industries and transport, construction and landfills influence the content of pollutants in soils (Meuser, 2010; Rodríguez-Seijo, et al., 2017); in terms of the sources of contamination of potential harmful elements, emissions resulting from vehicle traffic are mostly caused by wear of vehicular components like break lining, tire wear off and exhaust, as well as from incomplete fuel combustion, fuel additives or oil leaking from vehicles (Silva et al., 2021). The urban spaces are frequently located next to important busy avenues, surrounded by old dwellings and the urban garden may be found along motorways or arterial roads, railway lines and in prior industrial areas. Such sources of pollution can contaminate the soils of the urban spaces, mainly with potential harmful elements (As, Cd, Cr, Pb, Ni, Co, among others).

The aim of our study was to investigate element concentration in soil and *Citrus* sp. plant leaves along an urbanization gradient in Valencia, Spain. Our hypotheses are as follows:

(i) there are significant differences in pollutant levels in soils and *Citrus* trees among the stages of urbanization along the urbanization gradient (urban, periurban and rural area), (ii) the highest metal concentration in soil and leaves are in the urban, while the lowest concentration is in the rural area.

## Material and methods

### Study area

The soil and plant samples used for this study originate from municipalities in Valencia province, located in Eastern Spain. In the province of Valencia, the climate is Mediterranean; the average annual temperature was 17 ºC and the accumulated rainfall was 697 mm in 2022 (GVA, 2024). Valencia is the third most populated city in Spain with a population of approximately 809501as of 2023 (Cens de població, 2023). The area of the city is 135 km^2^ resulting in a population density of 5,662/km^2 (^Ayuntamiento de Valencia, 2024). Valencia is considered as a port city with an 1 international airport used for touristic and economic purposes less than 10 km from the city.

In the case of *Citrus sp*. in Valencia, approximately over 15,000 individual *Citrus* trees were reported by farmers in 2019 and Valencia is renowned for being the 2^nd^ largest orange exporter for the European Union (Torregrosa et al. 2019, Nyambura & Bonorchis 2022), this species is used as a crop tree as well as an ornamental tree in parks and gardens. The choosen *Citrus sp*. trees were healthy, and neither pests nor symptoms of disease were found by visual examination. The sampling areas were located in 5 urban areas (garden areas in the city of Valencia), 2 periurban areas (Valencia and Paiporta Cities), and 7 citrus ochard in rural areas of municipalities (Albal, Almusafes, Alzira, Guadasuar, Paterna, Picanya, Polinya del Xúquer) in the south of the Valencia city. We randomly chose 2-5 tree specimens at each sampling site, and we collected 20-30 leaves from each tree specimen. Samples were stored in paper bags and placed in the refrigerator at -21 °C. At each sampling area, soil sample, between 1-2 kg, was composed of 6 sub-samples, which were manually mixed with a small hand spade in a plastic container, and sifted to 2 mm in the field. All samples were taken from a depth of 0–20 cm with a Teflon-coated hand auger. Samples were placed into poly bags and were stored in at 4 °C in the refrigerator.

### Sample preparation and analyses

Leaves samples were dried in an oven at 50°C for 48 hours and pulverized. Directly after the homogenization step, 0.1000 g of the leaf samples measured on an analytical balance into 100 mL beakers with known mass. The beaker containing the leaf samples would then be placed in an oven at 105°C overnight to completely evaporate the water content. Following the drying process, the mass of the leaf sample was remeasured using an analytical balance to determine the exact mass of the sample. After this step, the leaf sample was ready for the digestion process.

Soil samples were air-dried. Soil pH was measured in soil:water suspension 1:5 v/v (UNE, 1999), and electrical conductivity (EC) was measured by suspension of soil in water at a ratio of 1:5 w/v (UNE, 2001), particle size distribution by Bouyoucos densimeter (MAPA, 1994). The equivalent calcium carbonate content (CaCO_3_) was determined by the volumetric calcimeter method (ISO 10693, 1995). Soil organic matter (SOM) was measured by oxidation with potassium dichromate in the presence of sulphuric acid and by subsequent titration with ammonium ferrous sulphate (Walkley & Black, 1934). Cation exchange capacity (CEC) and exchangeable sodium were determined by Ac-Na/Ac-NH_4_ method (MAPA, 1994). The exchangeable sodium percentage was calculated as follow:

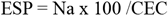

Relative to elements determination, 0.1000 g of the soil was directly measured into a 100 mL beaker of known mass directly after the collection step. The beakers were then similarly placed into an oven at 105°C overnight and on the following day, the mass of the soil samples were measured on an analytical balance. For the first step of the digestion process, both the soil and leaf samples were pre-treated with 5 mL of 65% HNO_3_ overnight. On the following day, the samples were placed onto a hot plate to completely evaporate the HNO_3_ solution. This was an important step as the digestion process requires the use of concentrated hydrogen peroxide (H_2_O_2_) which is able to react with the 65% HNO_3_ to form gaseous NO_X_ alongside an exothermic reaction releasing high amounts of heat. To decrease the likelihood of this incident, after evaporating the HNO_3_, 2 mL of deionized water was added into the beaker with the samples and allowed to evaporate completely on the hot plate. After this step, 200 μL of 30% (m/m) H_2_O_2_ was added into the beaker and once again placed onto the hot plate until complete evaporation. The last step for the digestion process was the addition of 10 mL of 1% HNO_3_ into the beakers and following this, the samples were ready for analysis using ICP-OES. Peach leaves (1547) CRM were used, and the recoveries were within 10% of the certified values for the elements (Simon et al. 2013, 2014, 2016, 2021).

### Statistical Analyses

Statistical analyses were conducted with SPSS Statistics 20 (IBM) statistical software. The normality of the distribution was tested using the Shapiro–Wilk test. The homogeneity of variances was tested with Levene’s test. The differences among samples were tested using analysis of variance (ANOVA) for each variable. Tukey’s test was used for pairwise comparison between the groups. Canonical discriminant analysis (CDA) was used to reduce dimensions and to identify those variables that most efficiently discriminated the study area as the dependent variable (De Sá, 2007).

## Results

Table 1 shows the mean values of the physical and chemical characteristics of the soils grouped in urban, peri-urban and rural areas. The texture of the rural and periurban soils was clay loam, very common in soils dedicated to *Citrus sp*. cultivation, however urban soils was loam, which is related to the fact that these soils are part of parks and gardens and are prepared to have a granulometric composition suitable for the development of garden plants. The pH of urban soils was slightly basic, while rural and periurban soils were moderately basic. The EC was low in the soils of all areas, with no salinity problems. The SOM and CaCO_3_ of rural and periurban soils were similar to conventional citrus crop soils of Valencia (Hondebrink et al., 2017). There were significant differences for the content of clay, pH, EC, SOM, CaCO_3_ and ESP (Table 1) (p < 0.05). Significantly lower results were found in the urban area for the content of clay and pH. While the highest results were also found in the urban area for EC, SOM, CaCO_3_ and ESP.

**Table 1.**
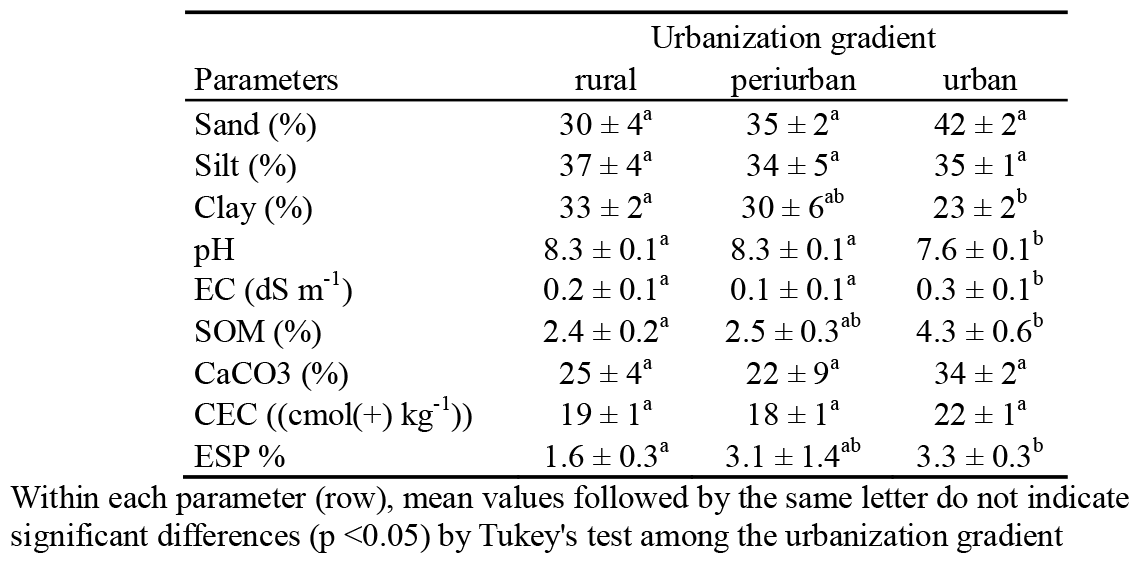
Main physical and chemical characteristics of soils in a urbanization gradient (mean ± SE). Notations: EC = electrical conductivity; SOM = soil organic matter; CEC = cationic exchangeable capacity; ESP = percentage of exchangeable sodium.

Table 2 shows the mean values of elemental concentration of the soils grouped in urban, peri-urban and rural areas. In relation to potentially toxic elements, urban soils showed higher average concentrations of Cr, Cu, Pb, Sr and Zn than periurban and rural soils, although these differences were not statistically significant. The other potentially toxic elements analyzed have very similar values among the different areas. The order of contents of potentially toxics elements were: Al>Sr>Zn>Pb>Cu>Cr>Ni>Cd>Co in rural soil, Al>Sr>Zn>Pb>Cr>Cu >Ni>Cd>Co in periurban soils, and Al>Sr>Zn>Pb>Cu>Cr >Ni>Cd>Co in urban.

**Table 2.**
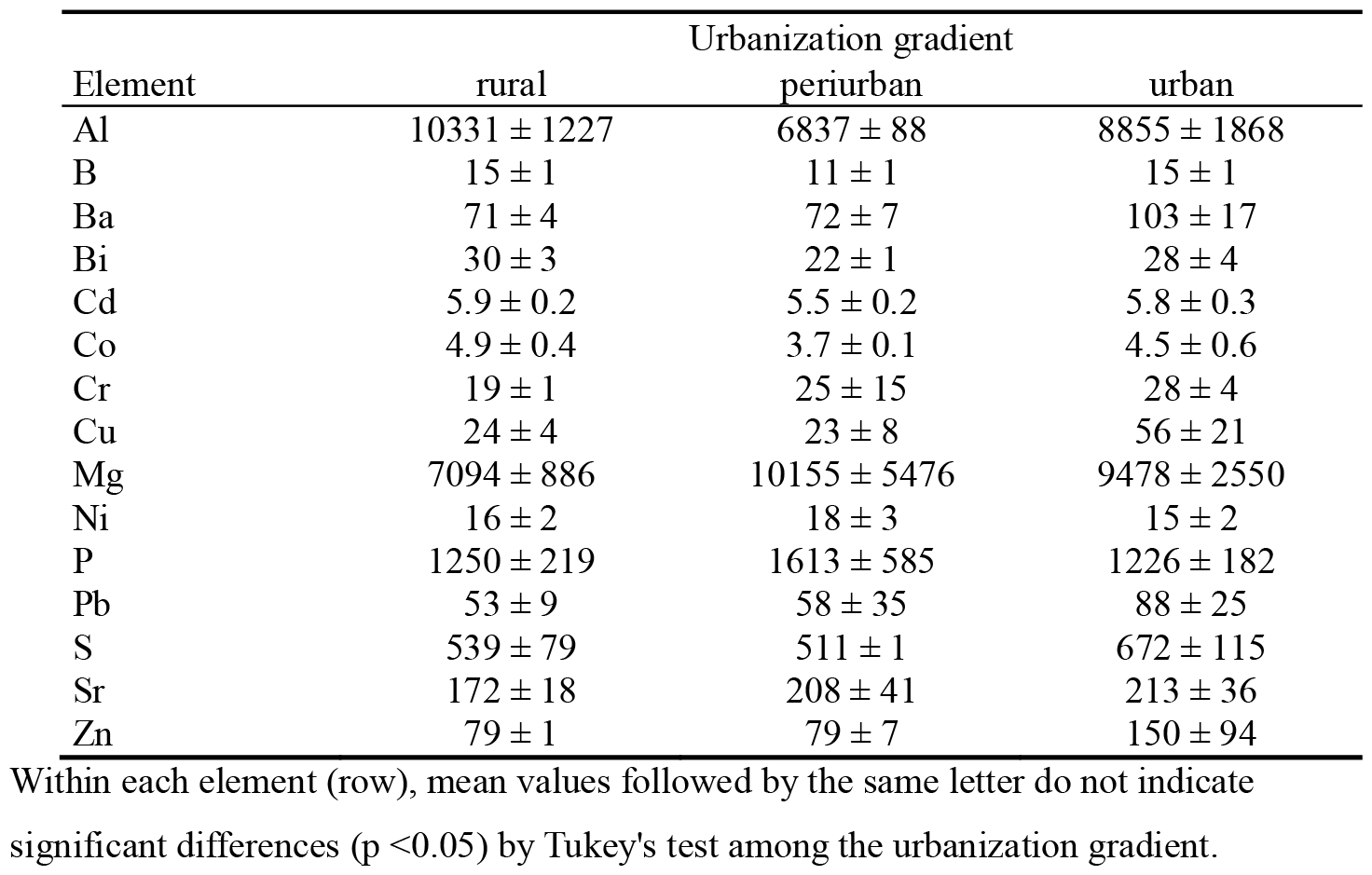
Elemental concentration (mg kg^-1^) of soils grouped in rural, periurban and urban(mean ± SE).

Based on main physical and chemical parameters of soil, canonical discriminant analysis (CDA) showed total separation among the studied areas (Fig. 1). The first discriminant functions (CDF1) contributed to 79.7% of the total variance, while the second one (CDF2) contributed 20.3% of the total variance. The canonical correlation was 0.963 for CDF1 and 0.873 for CDF2. Statistical differences were not found based on the CDF1 and CDF 2 (p = 0.057 and p = 0.260). Similar separation was found in the case of elemental concentration of soil (Fig. 2). The first discriminant functions (CDF1) contributed to 94.1% of the total variance, while the second one (CDF2) there was 5.9% of the total variance. The canonical correlation was 0.978 for CDF1 and 0.763 for CDF2. Statistical differences were also not found based on the CDF1 and CDF 2 (p = 0.162 and p = 0.773).

**Figure 1.**
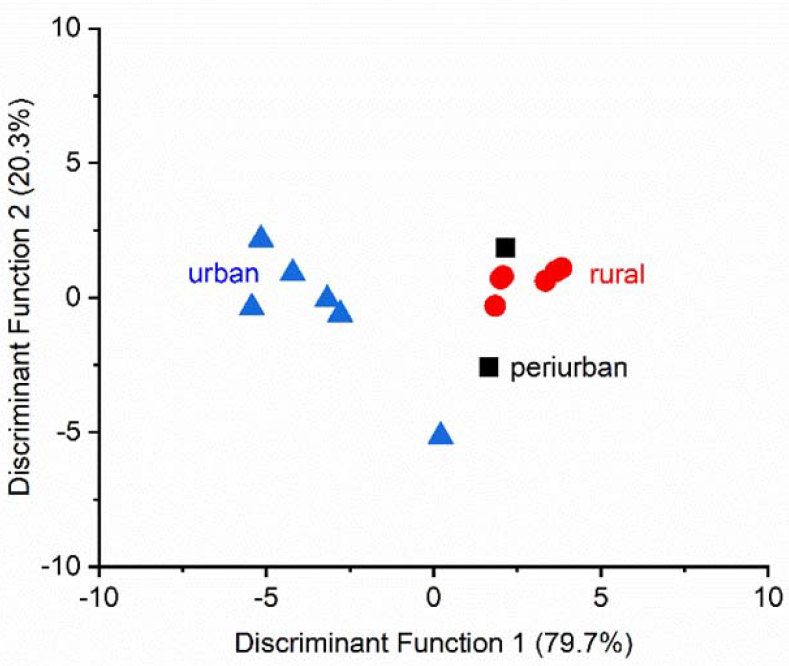
Scatter plot of canonical discriminant analysis based on the main physical and chemical parameters of soil among the studied areas.

**Figure 2.**
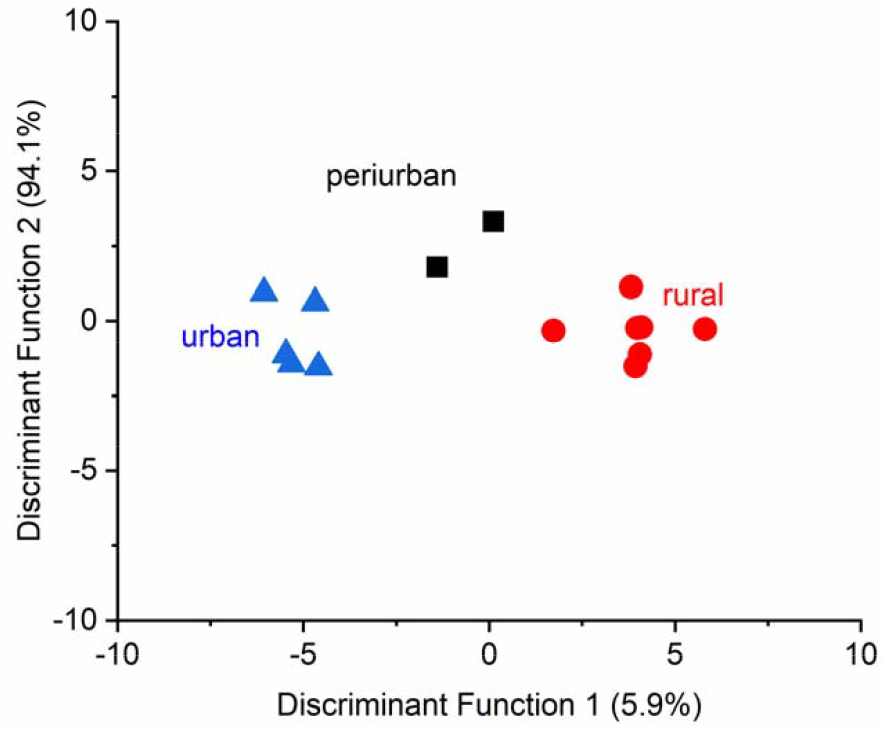
Scatter plot of canonical discriminant analysis based on the elemental concentration of soil among the studied areas.

The results of the elemental analysis of citrus leaves from the three areas studied are shown in table 3. The highest concentrations of Cr, Cu and Ni were obtained in the peri-urban area, while the rest of potentially toxic elements analyzed showed similar values between the peri-urban and urban areas, being higher than in the rural area. The ANOVA results show no significant differences in elemental concentration in *Citrus sp* leaves among urban, periurban and rural areas, except in the case of Cu (F = 5.397, p = 0.023), where periurban values were significantly higher than in rural (Table 3, Fig. 3)

**Table 3.** Elemental concentration (mg kg^-1^), of *Citrus sp*. leaves grouped in rural, periurban and urban (mean ± SE).

| Parameters | Urbanization gradient |  |  |
| --- | --- | --- | --- |
|  | rural | periurban | urban |
| Al | 62 ± 12 | 118 ± 79 | 97 ± 12 |
| B | 108 ± 26 | 65 ± 1 | 88 ± 26 |
| Ba | 18 ± 1 | 22 ± 6 | 22 ± 2 |
| Ca | 28477 ± 1436 | 27852 ± 2370 | 30569 ± 2047 |
| Cd | 2.7 ± 0.3 | 3.4 ± 0.1 | 3.7 ± 0.3 |
| Cr | 0.3 ± 0.1 | 1.6 ± 1.5 | 0.5 ± 0.1 |
| Mg | 3085 ± 292 | 3967 ± 714 | 2983 ± 111 |
| Ni | 5.3 ± 0.9 | 6.8 ± 0.6 | 4.9 ± 0.9 |
| P | 1268 ± 51 | 1193 ± 181 | 1179 ± 123 |
| Pb | 3.8 ± 0.5 | 4.8 ± 0.1 | 4.6 ± 0.7 |
| S | 2637 ± 127 | 2986 ± 40 | 3833 ± 540 |
| Sr | 406 ± 35 | 504 ± 37 | 501 ± 58 |
| Zn | 34 ± 1 | 39 ± 10 | 38 ± 3 |

Within each element (row), mean values followed by the same letter do not indicate significant differences ( $p < 0.05$ ) by Tukey's test among the urbanization gradient.

**Figure 3.**
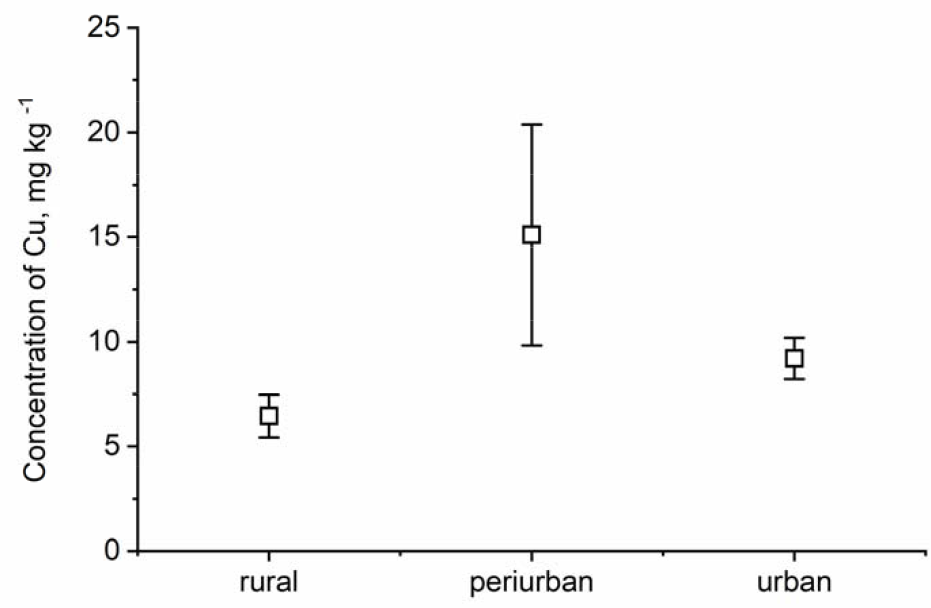
Elemental concentration of Cu (mean ± SE) among the studied areas.

**Figure 4.**
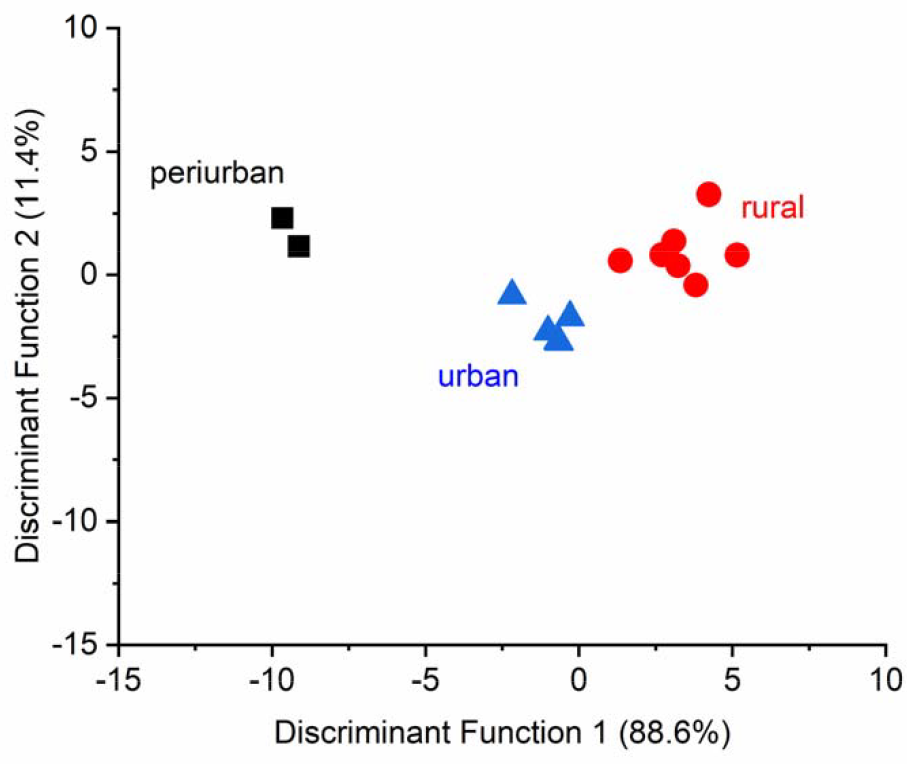
Scatter plot of canonical discriminant analysis based on the elemental concentration of *Citrus sp*. leaves among the studied areas.

Based on elemental concentration of *Citrus sp*., significant differences were not found among urban, suburban, and rural areas using canonical discriminant analysis (CDA), although total separation was found (Fig.4). The first discriminant functions (CDF1) contributed to 88.6% of the total variance, while the second one (CDF2) contributed 11.4% of the total variance. The canonical correlation was 0.980 for CDF1 and 0.868 for CDF2. Statistical differences were not found based on the CDF1 and CDF 2 (p = 0.188 and p = 0.590).).

## Discussion

Contrary to our hypotheses the main chemical and physical parameters and elemental concentration for the soil did not differ significantly in Valencia, Spain among the different areas along the urbanization gradient. Also, as hypothesized, we found differences among areas based on the Cu concentration of *Citrus* sp. leaves, which a is a typically anthropogenic element.

The absence of significant differences in the content of potentially toxic elements in the soils along the urbanization gradient, could be related to the fact that the rural soils studied, in our case are dedicated to citrus cultivation, although are not subjected to urbanization, soils are subjected to anthropogenic action due to the use of agrochemicals for crop production, which can contribute heavy metals to the soil, in this sense Gimeno-García et al. (1996) demonstrated that the use of agrochemicals contributes significant amounts of Cd, Cu, Pb and Zn to rice farming soils in Valencia. In our study, in general, no variations were observed in the order of the elements among urban, periurban and rural areas, which seems to indicate that the sources of contamination in each of the areas would contribute the same potentially toxic elements.

In the soils of rural areas, the mean value of Cd was higher, while Co and Cr were lower, than those obtained in agricultural soils of the Mediterranean area in Spain (Roca-Perez et al., 2010), on the other hand, Cu, Ni, Pb and Zn presented values similar to those obtained in the paper referred to previously. With regard to urban or peri-urban soils, Cd values are high for this type of soils (Sager, 2020), however similar values have been obtained in soils of the cities of Trondheim (Andersson et al., 2010), and Sopron (Horváth et al., 2012); In the case of Co, Cr, Cu Ni, Pb and Zn, the values obtained in this study are within the range of values for urban soils obtained in cities such as Salerno (Italy), Sporon (Hungary), Vigo (Spain), Mexico DC (Mexico), Vienna (Austria), etc (Sager, 2020). Similar to our findings, Francos et al. (2016) also measured the surface soil properties before and after rainfall and wildfire events in Catalonia. Within the comparison the data for the rural measurements was used as the majority of the data collected by Francos et al. (2016) were originating from areas susceptible towards wild fire such as forests. In terms of the major elements as calcium, potassium and phosphorus were measured by Francos et al. (2016) with the values of 30230 mg/kg for calcium, 338 mg/kg for potassium and 43.5 mg/kg for phosphorus. The percent difference between the measured calcium, potassium and phosphorus concentration were ±115%. ±165% and ±187% respectively. In terms of the minor non-toxic element. they measured a concentration of 401 mg/kg for sodium which was a percent difference of -20% with the measured sodium concentration.

In the case of Valencia leaf samples, the major elements were Ca as the most abundant followed by K, S and Mg. The explanation for the presence of S in the Valencia leaf sample was similarly attributed as the calcium found in the Valencia soil sample as the elements are added from the use of fertilizers. The use of S in citrus production increases the yield of the citrus tree by 23-57% (https://www.yara.us/crop-nutrition/citrus/). Additionally, the immense magnitude difference between the most abundant element and the following element can still be observed in the leaf samples of Valencia as the proportion between the concentration is approximately 3.88x between calcium and potassium. Observed pattern for copper concentration is highest in peri-urban sites followed by urban and rural sites in the Valencia leaf sample.

The values of Cr, Cu, Ni and Pb in leaf samples of *Citrus* sp from urban and periurban areas obtained in this study are lower than those reported for the same species in the city of Athens (Sawidis et al. 2012). This could be related to the lower degree of pollution in the city of Valencia compared to Athens during the time period in which the study was carried out. A study conducted in 2012 (Rodriguez Martín et al. 2015) in which leaves of trees from the botanical garden of Valencia were analyzed showed slightly lower levels of Pb and Zn, higher levels of Cr and similar values of Ni, Cu and Cd with respect to those obtained in this study in the urban area of the city of Valencia.

Marin et al. (2018) reported a decreasing tendency with the assessment of metal levels in foodstuffs in Valencia. While, Martin et al. (2013) demonstrated the anthropic effect on the soil based on the Cd, Pb and Zn concentration in Spain. They found spatial patterns with significant differences in territorial localisation. Ramos-Miras et al. (2020) found moderate ecological risk with the study of mercury and chromium in greenhouse soils in another region of Spain, in Almeria Valencia.

Rodríguez-Martín et al. (2015) reported that urban activities in Valencia have substantially raised levels of Cr, Ni and Cd in the urban atmosphere, which consequently increased atmospheric deposition due to changes in human activities over more than 70 years of urban growth. Lowering Pb concentrations and Zn stability agree with the control of leaded fuel combustion. Furthermore, no significant differences were detected in the concentrations of other metals. In short, metal deposition can be dangerous in the future if an increasing trend in Cr, Ni and Cd persists. At present however, the metallurgic industry, waste incineration, power generation and other anthropogenic sources now have the technology to reduce the atmospheric deposition of most elements to control pollution sources.

## Conclusion

Based on our findings Valencia did not exhibit a mathematical relationship between the element concentrations and the urban gradient except for the case of copper in the leaf samples.

Conclusively. this study assessed a range of elements from soil and leaves samples the pollution levels within an urban gradient using analytical methods and technologies. However, this research can be expanded on further by studying different urban cities to determine a larger or more concise pattern between element concentration and urban interactions in the context of pollution.

## Acknowledgements

Research was funded by the 2021 1 2 4 TÉT 2021 00049.

